# Synaptic function and sensory processing in ZDHHC9-associated neurodevelopmental disorder: a mechanistic account

**DOI:** 10.1101/2024.03.28.587155

**Authors:** Rebeca Ianov Vitanov, Jascha Achterberg, Danyal Akarca, Duncan E. Astle, Kate Baker

## Abstract

Loss-of-function *ZDHHC9* variants are associated with X-linked intellectual disability (XLID), rolandic epilepsy (RE) and developmental language difficulties. This study integrates human neurophysiological data with a computational model to identify a potential neural mechanism explaining *ZDHHC9*-associated differences in cortical function and cognition. Magnetoencephalography (MEG) data was collected during an auditory roving oddball paradigm from eight individuals with a *ZDHHC9* loss-of-function variant (ZDHHC9 group) and seven age-matched individuals without neurological or neurodevelopmental difficulties (control group). Evoked responses to auditory stimulation were larger in amplitude and showed a later peak latency in the ZDHHC9 group but demonstrated normal stimulus-specific properties. Magnetic mismatch negativity (mMMN) amplitude was also increased in the ZDHHC9 group, reflected by stronger neural activation during deviant processing relative to the standard. A recurrent neural network (RNN) model was trained to mimic recapitulate group-level auditory evoked responses, and subsequently perturbed to test the hypothesised impact of *ZDHHC9*-driven synaptic dysfunction on neural dynamics. Results of model perturbations showed that reducing inhibition levels by weakening inhibitory weights recapitulates the observed group differences in evoked responses. Stronger reductions in inhibition levels resulted in increased peak amplitude and peak latency of RNN prediction relative to the pre-perturbation predictions. Control experiments in which excitatory connections were strengthened by the same levels did not result in consistently stable activity or AEF-like RNN predictions. Together, these results suggest that reduced inhibition is a plausible mechanism by which loss of ZDHHC9 function alters cortical dynamics during sensory processing.

Graphical Abstract

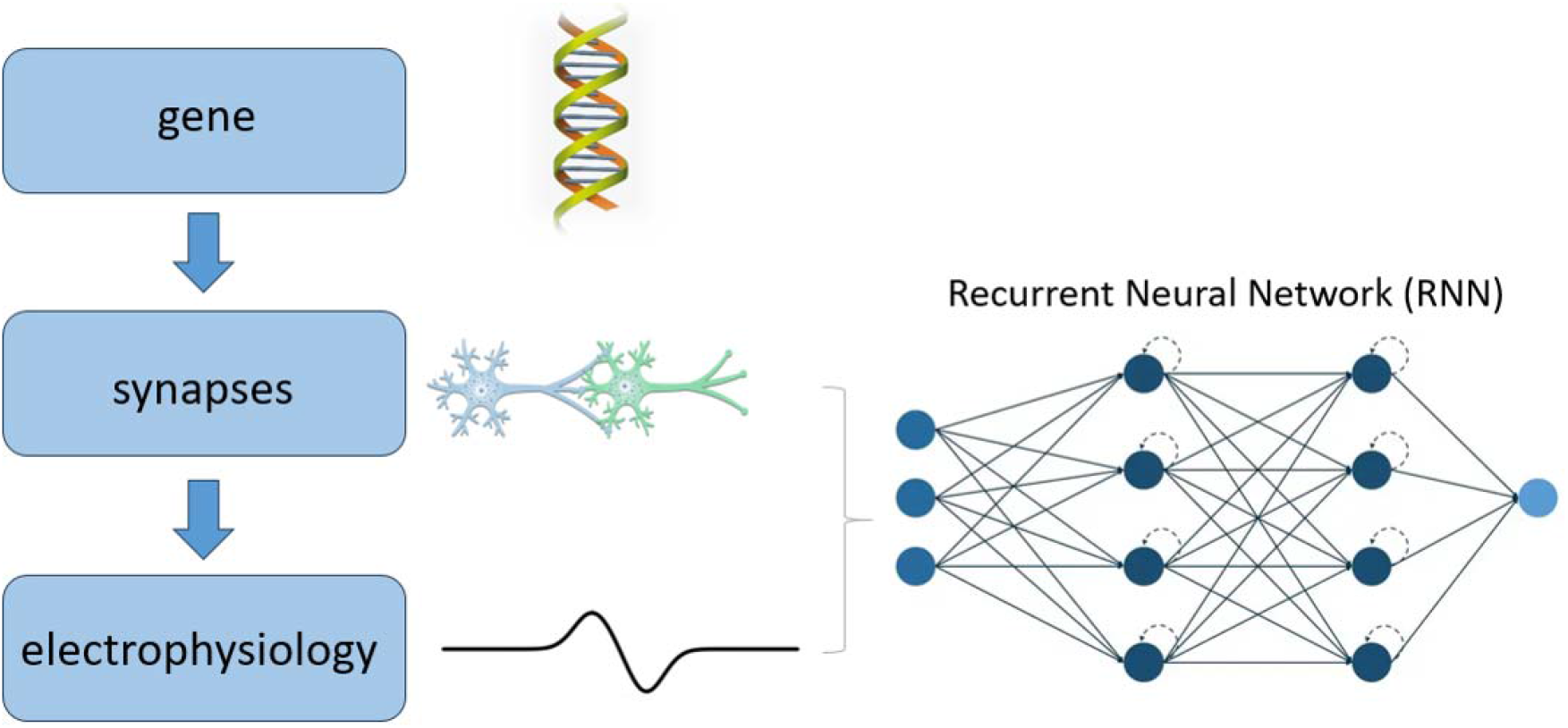

In the current study, we employed a bottom-up approach to study the impact of synaptic-level alterations associated with *ZDHHC9* variants on cortical function in healthy and *ZDHHC9*-deficient participants. To achieve this, a recurrent neural network model was developed to recapitulate MEG-derived auditory evoked responses and subsequently perturbed in order to determine effects on resulting dynamics. We show that reduced network inhibition recapitulates empirical observations, specifically increased response amplitudes, delayed peak latencies and increased mismatch negativity. These results offered a mechanistic account on the impact of *ZDHHC9*-associated synaptic alterations on auditory processing.

## Introduction

Cognition is sculpted during development by a myriad of genetic, cellular and systems-level mechanisms. Studying rare single gene disorders related to intellectual disability (ID), in combination with neural network models of cognition, could provide insights into specific mechanisms contributing to developmental cognitive difficulties. When made computationally tractable, specific cellular and systems-level mechanisms associated with genes of interest might offer explanatory support for the aetiology of cognitive impairment. Additionally, simplified computational models of the brain can be trained to perform tasks and then systematically perturbed to recreate ‘disorder-like’ model predictions. This framework is in line with the theoretical view that genes, brains, and artificial neural networks have similar underlying goals, which are to maximise the probabilities of achieving objectives, whether these be protein function, neural systems and cognitive functions, or good task performance, respectively (1). However, this approach has not previously been applied to rare genetic disorders, owing to the limited data availability across levels for these groups, particularly functional neuroimaging at appropriate temporal resolution. In this study, we trial the approach by employing a neural network model of auditory processing as a tool for mapping genetically-driven alterations to systems-level activity, in a group of individuals with cognitive impairment of known genetic origin.

A relevant gene for studying the emergence of intellectual disabilities is *ZDHHC9*, which encodes a palmitoylation enzyme (ZDHHC9) involved in the post-translational modification and intracellular trafficking of specific target substrates (2, 3). Loss of function *ZDHHC9* variants have been associated with mild to moderate ID (3), oromotor speech difficulties and language impairments (4), often coexisting with rolandic seizures (4, 5). Comorbidity between rolandic seizures, speech and language difficulties have been commonly observed in non-ZDHHC9 cohorts, but the mechanisms linking these symptoms remain elusive (6, 7). Hence, discovery of a rare monogenic cause of these associations may highlight specific, symptom-relevant neurobiological processes.

MRI studies of individuals with ZDHHC9-associated ID identified neuroanatomical differences that may increase the risk for epilepsy and cognitive impairments, such as reductions in cortical thickness and connectomic deviations (4, 8, 9). Another study of the same participant group employed resting-state magnetoencephalography (MEG), and revealed differences in state activation duration as well as dynamic connectivity across networks, with the extent of case-control differences correlating with *ZDHHC9* expression levels (10). While the studies outlined above have described the neurological, behavioural, neuroanatomical and global MEG characteristics of *ZDHHC9*-associated XLID, local MEG characteristics and causal links remain unexplored.

At the molecular and cellular level, experimental studies of *ZDHHC9* loss-of-function point toward developmental differences in synaptic structure and function. Targets of the ZDHHC9 enzyme include the GTPase Ras, which promotes dendrite outgrowth, as well as GTPase TC10, which supports inhibitory synapse formation (5). A study of the impact of *ZDHHC9* loss-of-function in primary rat hippocampal cultures revealed shorter and less complex dendritic arbours and an increase in the ratio of excitatory-to-inhibitory (E:I) synapses (5). Moreover, *ZDHHC9* knockout mice showed spontaneous high-frequency spiking activity potentially reflecting non-convulsive seizures (5). Another important target of ZDHHC9 is post-synaptic density protein 95 (PSD-95), a synaptic scaffolding protein that plays a key role in bidirectional synaptic plasticity, which is essential for learning and memory (11). In summary, there is emerging evidence that *ZDHHC9* variants alter properties of neuronal development and plasticity important for maintaining synaptic E:I balance and, ultimately, optimal neural function.

The current study aimed to link ZDHHC9-associated ID participants’ neurophysiology to previously reported cellular and synaptic differences, within a computational framework. MEG data were recorded from participants with *ZDHHC9* variants and control participants during a passive roving oddball paradigm, to enable assessment of auditory change detection via MEG mismatch negativity (mMMN) (12, 13). A roving protocol was designed for the study since this is an efficient method for observing mismatch responses and is relatively independent of basic stimulus properties. In addition, the roving oddball paradigm has been widely used in MEG studies of clinical groups with cognitive impairment (14, 15). Taking inspiration from previous studies integrating neural network modelling with electrophysiological data (16, 17), we employed a recurrent neural network to test a causal model relating *ZDHHC9*-related synaptic alterations to observed differences in group-level event-related fields.

## Materials and Methods

### Participants

Eight male participants age 9-41 with ZDHHC9-associated X-linked ID were recruited to the study (ZDHHC9 group). Mean estimated full scale IQ was 65 (standard deviation 6). Clinical and cognitive characteristics of this group have been previously described (4). Seven individually age-matched male comparison participants were recruited, free of neurological and psychiatric disorders (control group). Informed consent was obtained from each participant or their parent / consultee. Ethical approval for the study was granted by the Cambridge Central Research Ethics Committee (11/0330/EE).

### Data acquisition

All MEG datasets were collected on a 306-channel high-density whole-head VectorView MEG system (Elekta Neuromag, Helsinki), consisting of 102 magnetometers and 204 orthogonal planar gradiometers, located in a light magnetically shielded room. Data were sampled at 1 kHz and signals slower than 0.01 Hz were not recorded. A 3D digitizer (FASTRACK; Polhemus) was used to record the positions of five head position indicator (HPI) coils and 50–100 additional points evenly distributed over the scalp, all relative to the nasion and left and right preauricular points. An electrode was attached to each wrist to measure the pulse and bipolar electrodes to obtain horizontal (HEOGs) and vertical (VEOGs) electrooculograms. Head position was monitored throughout the recording using the HPI coils.

### Stimuli and design

The experimental design was based on a roving oddball paradigm described by Cowan et al (12). This experimental scheme involved the repeated presentation (3-12 times) of standard stimuli of a particular frequency (250Hz, 500Hz, or 1000Hz) followed by a deviant tone of a different frequency which, in turn, is repeated and becomes the new standard (Figure 1). Tones were 50ms in duration and the inter-tone interval was fixed at 500ms (550ms stimulus onset asynchrony). Participants were instructed to watch a silent movie and ignore the tones.

**Figure 1:**
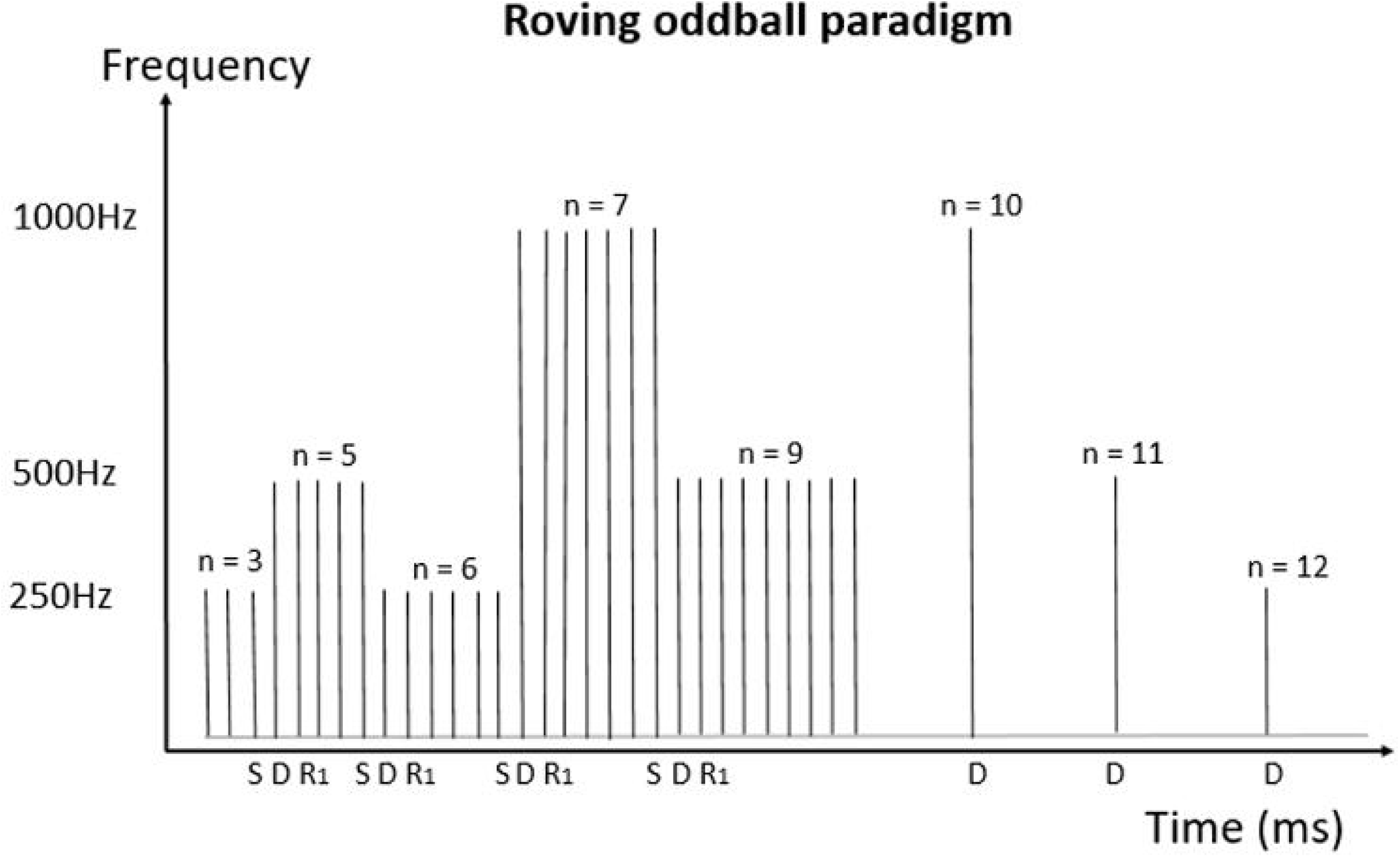
Roving oddball paradigm. A standard tone (S) of either 250Hz, 500Hz, or 1000Hz is presented randomly between 3-12 times. After this sequence of repetitions, the frequency of the tone changes (deviant tone, D), which then becomes the new standard through repetitions.

### MEG data pre-processing

The raw MEG data was pre-processed with the MNE package (version 1.0.3) (18) in Python 3.10. External noise was removed using a signal-space separation method and adjustments in head position within the recording were compensated for using Maxwell filtering. A sensor-space temporal independent components analysis (ICA) was used to automatically remove artefacts arising from blinks, saccades and pulse-related cardiac artefacts, and the outputs were manually checked by visual inspection. Data were epoched to a time window defined as 400ms pre-stimulus and 550ms post-stimulus to ensure all relevant event-related changes were contained within the epoch time window, and then down-sampled to 250Hz, baseline corrected and low-pass filtered at 30Hz utilising a Butterworth filter. All trials with larger peak-to-peak amplitude than 4e-12 Tesla (4000 fT) and smaller peak-to-peak amplitude than 1fT at each magnetometer were removed. For each participant, data were averaged across all trials to form the time-domain signals. Event-related field (ERF) analyses were also performed with the MNE package (version 1.0.3) (19).

### Data analysis

Auditory evoked fields (AEFs) across all stimuli types were computed for both groups. The grand average of all trial types was used to compute AEF time-domain signals. Peak amplitude and peak latencies of AEFs were automatically calculated at the magnetometer where the largest amplitude signal was detected. The precise latencies of M100 responses are useful indicators of the temporality of auditory processing (20). To account for a consistent 50ms delay in the tone stimulus presentation relative to the trigger timing in the scanner setup, a peak of activity at approximately 150ms post-trigger was automatically detected as equivalent to the widely reported M100 response (21). Differences in activity elicited by standard and deviant stimuli were analysed for both groups using non-parametric cluster-based permutation testing (22), which detects spatial clusters of sensor locations for which significant differences between trial types, within a certain time period, are found. All deviant responses (D) were compared to responses elicited by their preceding standard stimuli (S). A significance threshold of 0.05 (alpha) was used. mMMN was calculated as the mean absolute error (MAE) between the standard and deviant-evoked responses at the group level. Individual MMNs were calculated at the largest cluster significant at the group level.

### Neural network modelling

A recurrent neural network (RNN) consisting of an input layer, four hidden layers and an output layer was employed as a model of the auditory cortex. RNNs are a class of artificial neural networks where the connections between nodes can create a cycle, allowing output from some nodes to be fed as input to the same nodes. These recurrent connections allow RNNs to learn from sequences of inputs and are loosely analogous to feedback connections in biological neural networks. Each RNN node is analogous to a population of neurons, which can emit excitatory or inhibitory connections (weights). The model used is a discrete-time RNN, which processed the input at each timestep according to the recurrence formula (Equation 1), where the matrix W_hh_ captures the recurrent connections, h_t-1_ is the previous hidden states vector, W_xh_ is the feedforward weight matrix and the x_t_ is the input vector at timestep t. The RNN output at each timestep is obtained according to Equation 2, where W_hy_ is the weight matrix between the last hidden layer and the output units and h_t_ is the hidden state vector at the current timestep.

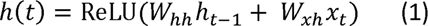

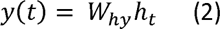

The RNN was trained in a supervised fashion, and the labels (targets) for the S and D inputs that were given to the RNN were obtained by adding Gaussian white noise to the control group post-stimulus S and D AEFs, respectively. 1200 targets were obtained for S and D responses, respectively. Before feeding the labels into the RNN model, the data was robust-scaled and flipped so that most values are above 0. Robust scaling sets the median and interquartile range to 0 and 1, respectively, and therefore maintains the directionality of the amplitude difference between S and D trial type AEFs (i.e the RNN predictions when D inputs are given have a higher amplitude than the S predictions, as observed empirically).

The RNN inputs were sound waveforms produced with a sampling frequency of 5000Hz. Standard inputs consisted of 3 sinewaves of the same frequency and deviant inputs consisted of the first 2 sinewaves of the same frequency and the third one of a different frequency. The frequencies used for the sinewaves were 250Hz, 500Hz and 1000Hz (Figure 1). These were converted into the time-frequency domain using the short-time Fourier transform (STFT), producing a representation of cortical input from the ascending auditory pathway (23). The STFT was performed on Hann-windowed segments of 125 samples with an overlap of 105 samples. Input features were fed through the model and its parameters (weights and biases) were optimised to minimise mean-squared-error (MSE) loss between model outputs at the current timestep, f_t_, and target evoked responses, or observed values at the current timestep, y_t_ as shown in Equation 3.

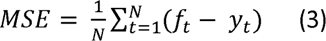

Adaptive moment estimation (Adam) optimization was used with a learning rate of .0002 and a dropout regularisation of 0.15 was used in the hidden layers. Each hidden layer had a rectified linear unit (ReLU) activation, whereas the final layer had a linear activation function. Connection weights between layers were initialised from a Glorot uniform distribution and recurrent weights were initialised as an orthogonal matrix from a normal distribution (24, 25).

A 70%:15%:15% train:validation:test split was used for training, tuning the model parameters (validation) and testing the model performance. The stability of RNN predictions in the three experiments was validated using principal component analysis (PCA). The units’ activations of the fourth hidden layer, represented by a matrix of size (64 [units], 138 [timepoints]), were transformed into principal component spaces of size (2 [components], 138 [timepoints]). This transformation preserves as much variance as possible from data in the original matrix while compressing them into fewer columns. The resulting neural latent space, obtained after PCA, is shown on a cartesian plane in Figure S4, which indicates the activity in the last hidden layer over the timecourse of the RNN prediction.

## Results

### Empirical MEG results

AEFs across all stimuli types were computed for the ZDHHC9 and control groups (Supplementary Figure S1). Mean peak activity occurred at 144ms post-stimulus in the control group, reaching a peak absolute distribution of 140fT, which corresponded to an expected M100 response. For the ZDHHC9 group, mean peak activity over this window occurred at 188ms post-trigger, reaching a maximal absolute magnetic distribution at 203fT – 44ms later in latency (p-value = 0.08) and 1.13X greater in magnitude (p-value = 0.3) than control subjects. The topographical plots computed at the peak latency (Supplementary Figure S1, right panel), indicated a clear dipolar pattern in both the left and right hemispheres in both groups.

Trial responses were then separated by frequency into 250Hz, 500Hz and 1000Hz stimulus responses for both the control and ZDHHC9 group, to explore whether there was a systematic relationship between the frequency of the stimulus and the latency of the AEF. Given the spatial tonotopic organisation of the auditory cortex (26), groups of neurons that respond to higher frequency stimuli are activated before those that respond to lower frequency stimuli. Thus, it was expected that higher frequency stimuli would result in shorter latency responses (26, 27, 28). In control participants, 250Hz, 500Hz and 1000Hz stimuli evoked an AEF with a peak at 156ms, 140ms and 132ms – decreasing in latency respectively as expected (Supplementary Figure S2). In contrast, the ZDHHC9 group showed delayed peak amplitudes and larger response amplitudes for the 500Hz and 1000Hz stimuli. In this group, the frequency-latency dependence was present for the 250Hz (latency: 0.192) and 500Hz stimuli (latency: 0.164), but the 1000Hz stimulus resulted in a delayed response (0.200ms). The two peaks present for the 1000Hz stimuli were due to wider inter-subject variability for the ZDHHC9 group. For all frequency stimuli, the responses in this group reflected a prolonged activation (Supplementary Figure S2).

Next, responses to all deviants (D) were computed and compared to their preceding stimulus (S) to provide an index of the mMMN response across all standard train lengths, for the control and ZDHHC9 groups separately (Figure 2). Non-parametric cluster-based permutation testing revealed statistically significant differences between S and D trials (P_corrected_ = 0.0008) in the control group at a single negative cluster in the right hemisphere, occurring over 12 channels, from 150-180ms after stimulus onset (Figure 2a). In the ZDHHC9 group, six significant clusters were identified i.e. mismatch responses were topographically more extensive and of higher mean peak amplitude in the case group (Figure 2b). The responses of the two groups could only be directly compared by calculating the average evoked responses at the eight sensors where significant S-D contrasts were found in both groups (Supplementary Figure 3). All averaged responses were significant within a time-frame of an expected mMMN, around 100ms after the end of stimulation (highlighted in yellow). Individual-level responses were computed for and averaged across these eight channels - differences in peak amplitudes between the two groups did not reach statistical significance for the standard-evoked responses (p-value = 0.09) or the deviant-evoked responses (p-value= 0.1).

**Figure 2:**
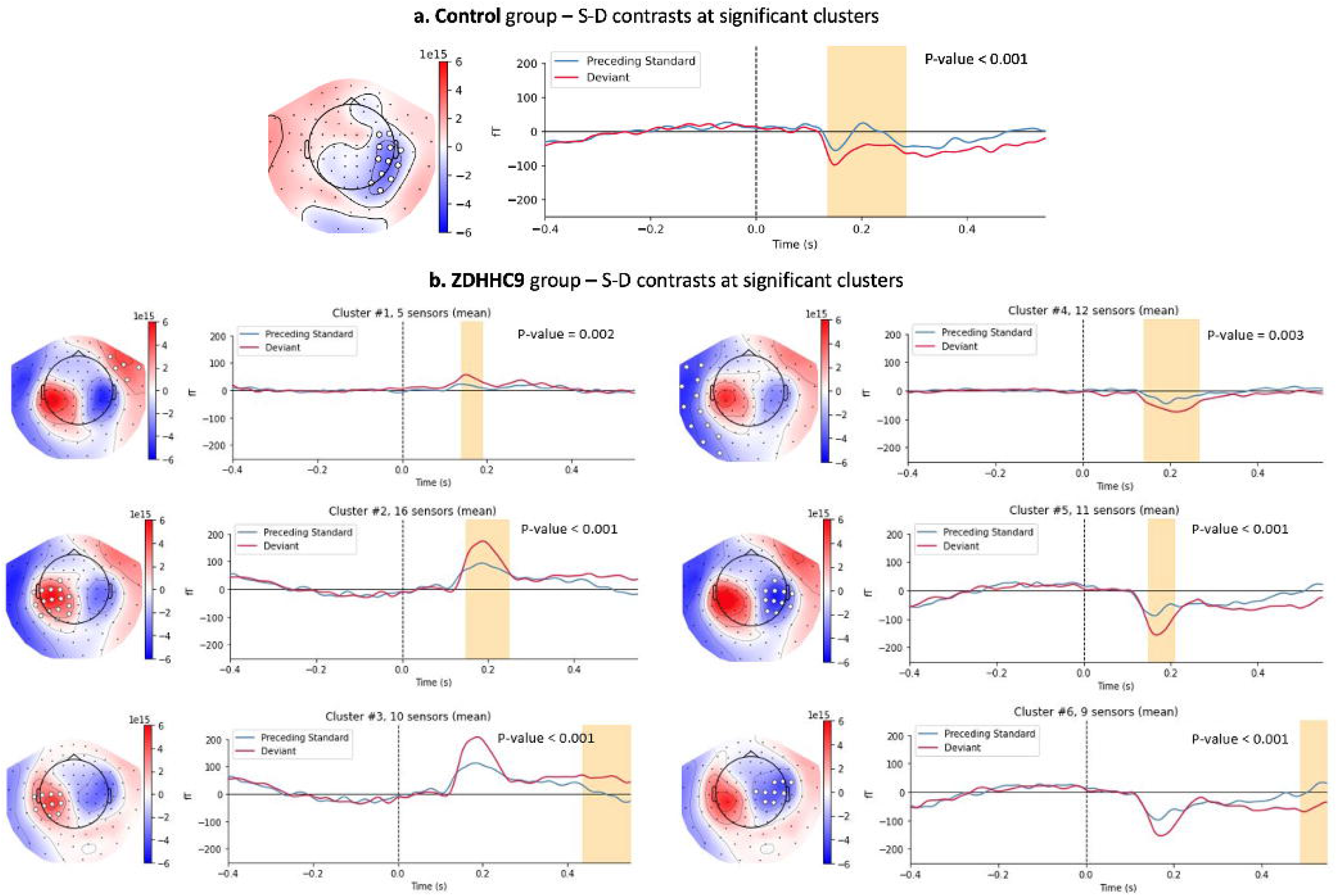
Cluster-based permutation testing. Permutation-based clusters of significant difference between standard-evoked and deviant-evoked AEFs in control and ZDHHC9 groups.

### Recurrent neural network model of neural dynamics in auditory processing

Figure 3 outlines the modelling workflow. Spectrograms representing standard tones and deviant tones were given to the RNN model as inputs and empirically-derived control group-level AEFs, on which Gaussian noise were added, served as corresponding labels. During training (10 epochs), MSE was minimised between the RNN predictions and the labels. Average model outputs over all standard and deviant types, after training, are shown in Figure 3e.

**Figure 3:**
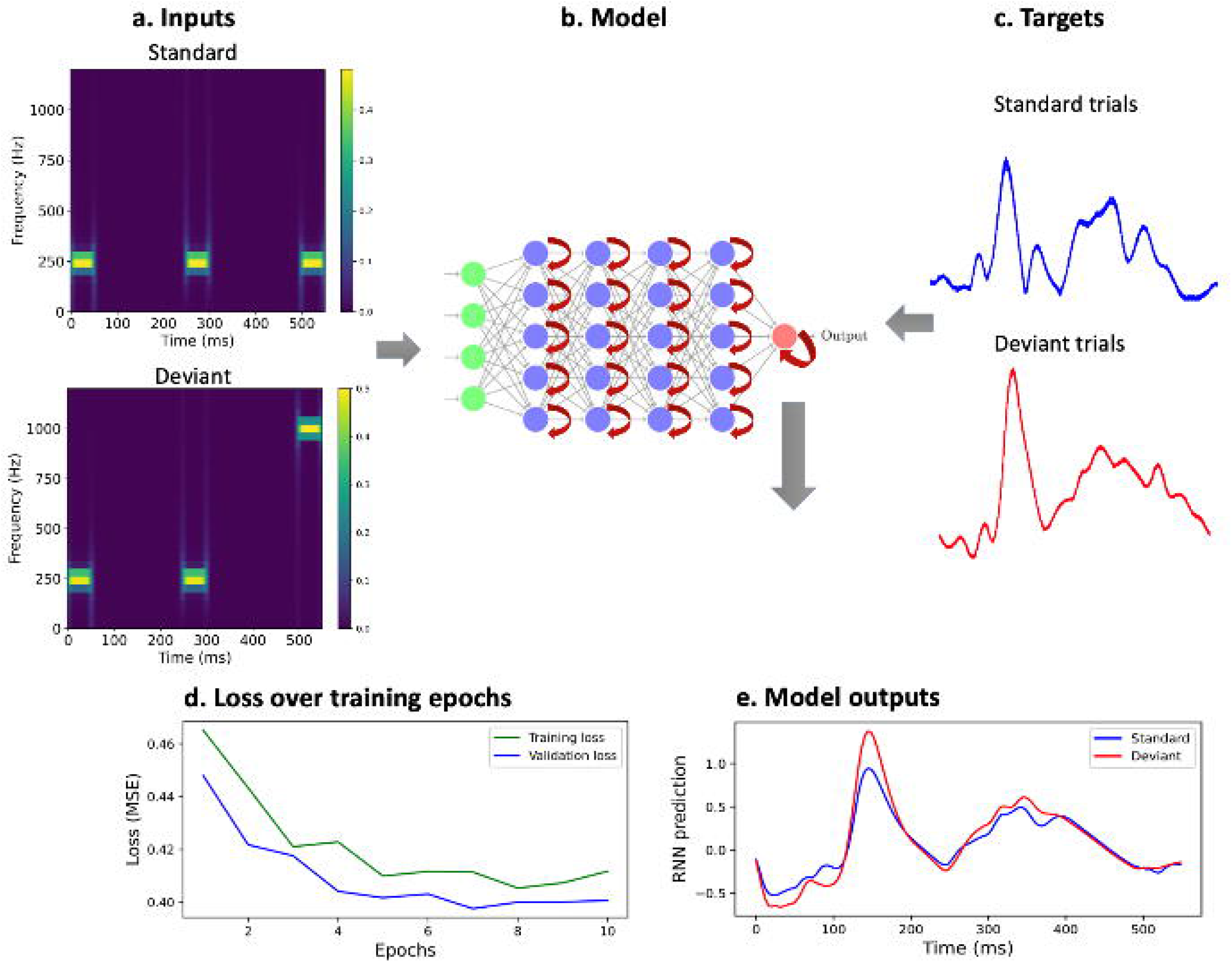
Overview of modelling workflow. a. Spectrograms of an example standard and deviant input used for the RNN. Three frequencies were used as in the roving oddball paradigm: 250Hz, 500Hz and 1000Hz. Deviant inputs had the first 2 tones of the same frequency and the third tone of a different frequency. The train set consisted of standard inputs of 250Hz and 500Hz, as well as the following sequences of tones, of which the third represented the deviant: 250-250-500 (Hz), 250-250-1000 (Hz), 500-500-1000 (Hz). The test set consisted of standard inputs of 1000Hz and tone triads including: 500-500-250 (Hz), 1000-1000-250 (Hz) and 1000-1000-500 (Hz). b. Simplified diagram of the hierarchical RNN architecture. Input layer (green) had 63 recurrent units, each hidden layer (4) had 64 units and the output layer had 1 recurrent unit. c. Targets were 1200 simulated AEFs obtained by adding Gaussian white noise (standard deviation 0.6) to the control group level post-stimulus AEF in response to standard tones. The same was done for deviant AEFs, resulting in 1200 simulated deviant AEFs. d. The RNN was trained for 10 epochs (i.e. iterations through the entire training dataset). The lower validation loss reflects the absence of dropout regularisation during validation, as opposed to training. e. RNN predictions to S and D inputs.

To understand how the hierarchy of the RNN’s layers corresponded to the types of inputs, we computed the RNN hidden layer activations for S and D inputs (Figure 4). This enabled us to qualitatively assess whether the network responded differentially based on the input and, if so, where in the network this was occurring. We found that network activity became gradually more diffuse in time from the first to the fourth hidden layers. In the first hidden layer, activations for the S and D inputs were largely similar, whereas in later layers of the hierarchy, the activity elicited by the D inputs became larger than that elicited by the S inputs. (Figure 4a, b).

**Figure 4:**
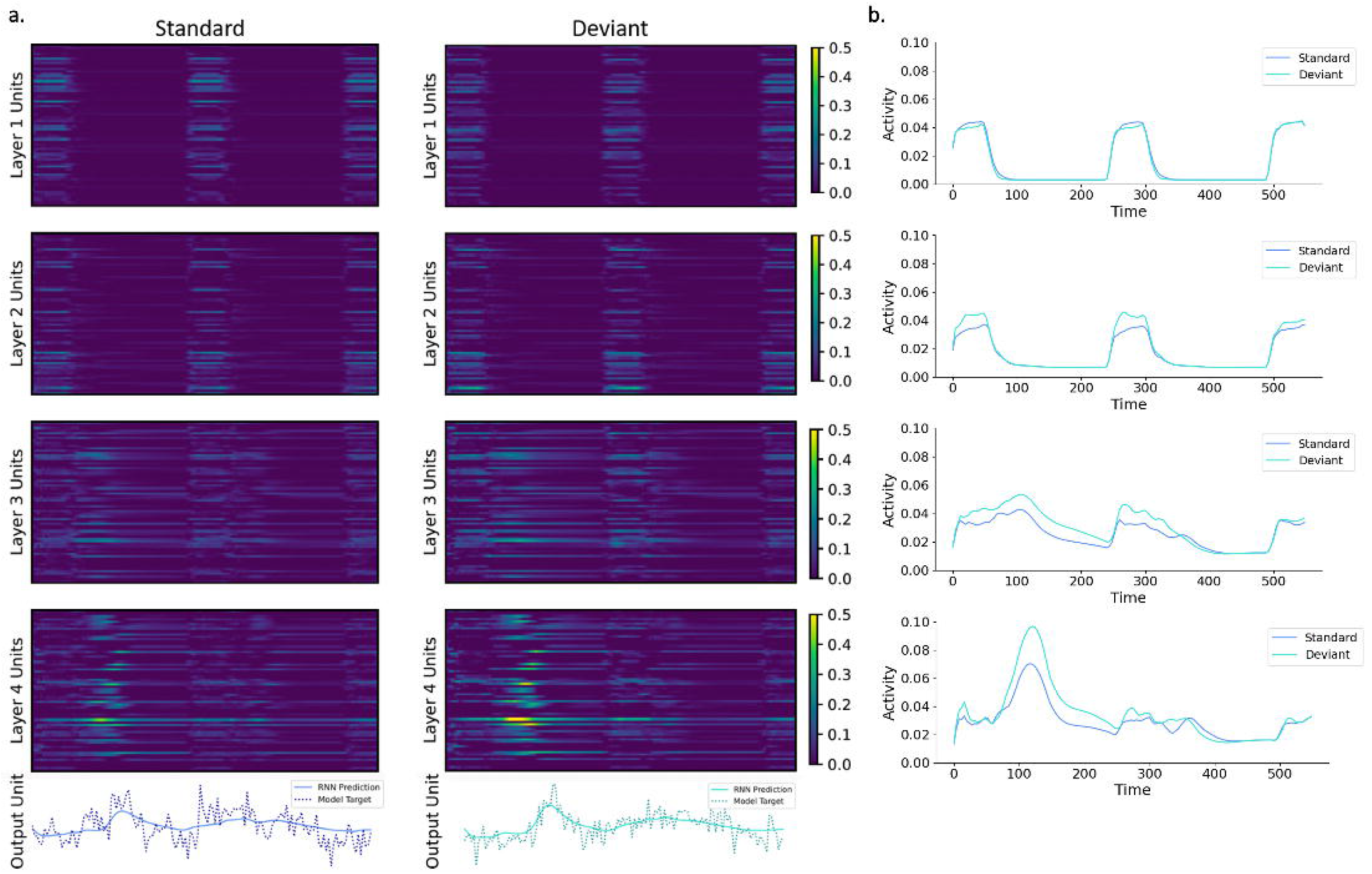
Hidden layer activations and output unit predictions for standard and deviant inputs. a. Model predictions and hidden layer activations plotted by layer (rows) and input condition (columns). The activations were computed as an average across all S input types and all D input types, respectively. The x-axis represents time. Amplitudes of hidden unit activations, shown in the four upper rows, increase towards the output. Patterns of hidden unit activations also spread out and become more complex with increasing layer depth. Visible differences in hidden unit response magnitudes between input conditions were also found. Model outputs and corresponding grand-average ERFs (with Gaussian white noise; s.d. = 0.6) are plotted in the bottom row. All panels display data from the end of stimulus onset (0ms) to 544ms after stimulus onset. b. Hidden layer (1–4) activations over time, averaged across the 64 hidden units.

### Reducing inhibition recapitulates auditory dynamics in the ZDHHC9 group

To test the impact of alterations mimicking the *ZDHHC9* loss-of-function phenotype (i.e. reduced inhibition) on the RNN output, we conducted a perturbation experiment by systematically reducing inhibition levels after network training and observed the effects on the network’s predictions.

In experiment 1 (“negative weight perturbation”, Figure 5a), the negative recurrent weights of the four hidden layers were reduced, in terms of absolute values, by eight arbitrary levels that resulted in stable changes in RNN output from the baseline: 0.5%, 1%, 1.5%, 2%, 2.5%, 3%, 3.5%, 4%. These alterations thus altered the average excitation-inhibition synapse ratio of ∼1:1 obtained after training. Two additional control experiments were performed, in which the outcomes were assessed of increasing excitation or concomitantly increasing excitation and reducing inhibition, for the same levels as in the first experiment. In experiment 2 (“positive weight perturbation”, Figure 5b), the positive recurrent weights were relatively increased by 0.5-4% (with the same increments as in experiment 1). In experiment 3 (“random weight perturbation”, Figure 5c), a random set of recurrent weights, including both positive and negative connections, were altered by the same levels (positive weights were increased and the absolute value of negative weights was decreased).

**Figure 5:**
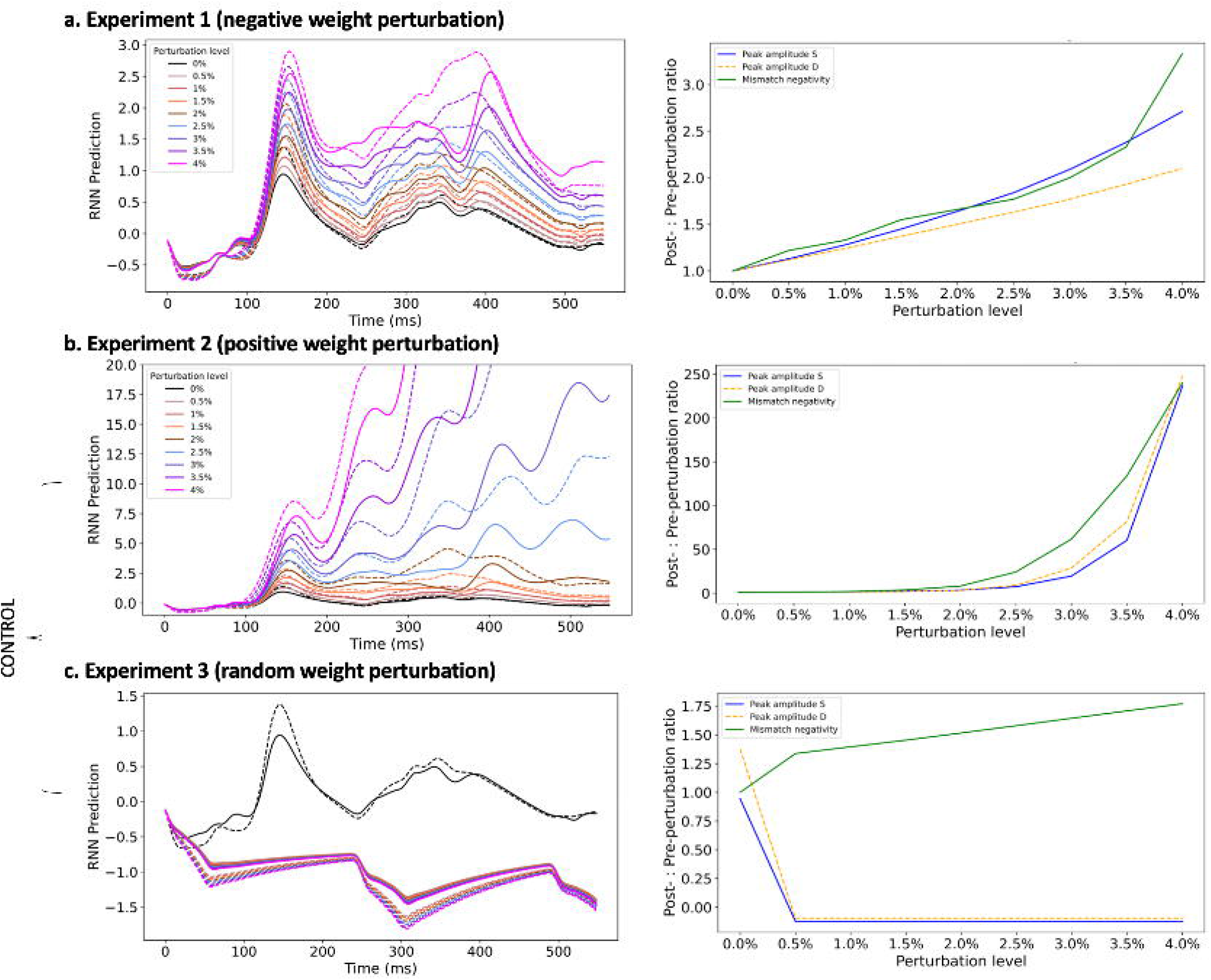
RNN predictions before and after each perturbation experiment. Predicted AEFs for standard and deviant inputs (left panel) and relative increases from initial RNN predictions (pre-perturbation) after each perturbation experiment (right panel). a. Experiment 1 (negative weight perturbation), b. Experiment 2 (positive weight perturbation) and c. Experiment 3 (random weight perturbation).

To quantify how well the model was fitting to the data, we quantified the value of the loss function, which usually decreases with improved data fitting (i.e. smaller differences between network predictions and labels). The loss values evaluated on the test set, which implied both S and D inputs, are shown in Table 1. Reducing inhibition (Experiment 1) resulted in the smallest MSE values up to 3.5% inhibition reduction, with small MSE increases to the next perturbation level, whereas increasing excitation levels led to unstable RNN predictions reflecting runaway excitation (Table 1). Experiment 3 resulted in relatively stable MSE values across the perturbation levels (Table 1). These results showed that inhibition reduction experiments best fitted the evoked response data, with small increases in perturbation strength resulting in small increases in loss values, consistent with our predictions.

**Table 1:** RNN mean squared error (MSE) values during test set evaluation for each experiment, across the perturbation levels.

| Test loss function (MSE) values |  |  |  |  |  |  |  |  |
| --- | --- | --- | --- | --- | --- | --- | --- | --- |
| Pre-perturbation: 0.4 |  |  |  |  |  |  |  |  |
| Perturbation level: |  |  |  |  |  |  |  |  |
|  | 0.5% | 1% | 1.5% | 2% | 2.5% | 3% | 3.5% | 4% |
| Experiment 1 | 0.39 | 0.43 | 0.50 | 0.63 | 0.87 | 1.28 | 1.93 | 3.21 |
| Experiment 2 | 0.44 | 0.71 | 1.72 | 6.31 | 53.23 | 368.7 | 2279.2 | 14323.8 |
| Experiment 3 | 2.19 | 2.22 | 2.26 | 2.30 | 2.34 | 2.34 | 2.42 | 2.46 |

We next analysed the model’s qualitative dynamical outputs under perturbation for the three experiments. The output unit predictions for S and D inputs were computed in each experiment and compared to baseline levels, pre-perturbation (Figure 5). Inhibition reduction (Experiment 1) resulted in predicted AEFs with linearly increasing amplitudes (Figure 5, Table 2) relative to the baseline levels, which mirrors the trend observed empirically between the control and ZDHHC9 groups (Figure 2). MMN showed nonlinear increases, especially at the highest perturbation levels. AEF peak latencies also increased from baseline, by 4ms, and remained constant across the 0.5-2.5% perturbations before a further 4ms increase at the 3-4% perturbations (Table 2). No differences in peak latencies were obtained between S trial predictions and D trial predictions (Table 2). Increased excitation (Experiment 2) resulted in exponential increases in AEF peak amplitude from the 4^th^ perturbation level onwards (Figure 5), accompanied by peak latencies in the latter half of the AEF window (Table 2). Perturbing a random set of weights (Experiment 3) resulted in opposite polarity AEFs with peak amplitudes and MMN varying minimally across the perturbation levels, and a constant peak latency at 300ms (Figure 5, Table 2).

**Table 2:** AEF metrics before and after perturbations.

a. Pre-perturbation AEFs
| Trial type | Peak amplitude | Peak latency | MMN |
| --- | --- | --- | --- |
| S | 0.94 | 140 | 0.09 |
| D | 1.37 | 140 |  |

b. Post-perturbation AEFs
| Experiment |  | Trial type | Perturbation level |  |  |  |  |  |  |  |
| --- | --- | --- | --- | --- | --- | --- | --- | --- | --- | --- |
|  |  |  | 0.5% | 1% | 1.5% | 2% | 2.5% | 3% | 3.5% | 4% |
| 1 | Peak amplitude | S | 1.07 | 1.21 | 1.37 | 1.55 | 1.74 | 1.98 | 2.25 | 2.56 |
|  |  | D | 1.54 | 1.71 | 1.88 | 2.07 | 2.25 | 2.43 | 2.65 | 2.89 |
|  | Peak latency | S | 144 | 144 | 144 | 144 | 144 | 148 | 148 | 148 |
|  |  | D | 144 | 144 | 144 | 144 | 144 | 148 | 148 | 148 |
|  | MMN |  | 0.11 | 0.12 | 0.14 | 0.15 | 0.16 | 0.17 | 0.2 | 0.3 |
| 2 | Peak amplitude | S | 1.26 | 1.65 | 2.12 | 3.30 | 6.97 | 18.48 | 57.1 | 223.4 |
|  |  | D | 1.81 | 2.3 | 2.8 | 4.5 | 12.4 | 39.65 | 112.7 | 341.8 |
|  | Peak latency | S | 144 | 144 | 148 | 400 | 500 | 508 | 544 | 544 |
|  |  | D | 144 | 144 | 144 | 344 | 516 | 544 | 544 | 544 |
|  | MMN |  | 0.13 | 0.18 | 0.35 | 0.77 | 2.3 | 5.87 | 12.7 | 22.8 |
| 3 | Peak amplitude | S | -1.38 | -1.39 | -1.41 | -1.42 | -1.43 | -1.44 | -1.46 | -1.48 |
|  |  | D | -1.61 | -1.64 | -1.67 | -1.69 | -1.72 | -1.75 | -1.78 | -1.8 |
|  | Peak latency | S | 300 | 300 | 300 | 300 | 300 | 300 | 300 | 300 |
|  |  | D | 300 | 300 | 300 | 300 | 300 | 300 | 300 | 300 |
|  | MMN |  | 0.12 | 0.13 | 0.13 | 0.14 | 0.15 | 0.15 | 0.16 | 0.17 |

### Validating the effect of decreased inhibition on latent dynamics aligning with the ZDHHC9 phenotype

Principal component analysis was performed on the post-perturbation activations of the 4th hidden layer after model evaluation on the test set (Figure S4). This provided a visualisation of the internal representation of the RNN just before the outputs are read by the output unit and are in line with the results in Table 2. Experiment 1 resulted in stable predictions across the perturbation levels, as activation in latent space at the first prediction timepoint was in proximity of activation level at the last prediction timepoint, as shown in the latent activity depicted in Figure S4a. The area covered by the latent space activity increased with the perturbation levels, reflecting the AEF amplitude increases (Figure S4a). The latent activity in Experiment 2, starting from the 5th perturbation level (2.5%) onwards resulted in values that maximally differed between the initial and last timepoints, highlighting runaway activity (Figure S4b). Experiment 3 resulted in different internal representations which remained stable across the perturbations (Figure S4c).

## Discussion

This study tested the hypothesis that empirical neurophysiological differences between a monogenic neurodevelopmental disorder group (*ZDHHC9*-associated ID) and control group are compatible with reduced inhibition in a network model of auditory processing. We observed that reduction in inhibition levels within an RNN model resulted in increasing peak amplitudes of model outputs, which qualitatively matched the case-control results. In contrast, increasing excitation or perturbing a random set of connections resulted in a phase shift of the RNN output, inconsistent with empirically-derived results.

Empirical MEG analyses focused on magnetic mismatch negativity to determine whether *ZDHHC9* variants are associated with differences in adaptive auditory processing. mMMN, an auditory evoked field reflecting the difference between the brain response to standard and deviant stimuli, captures auditory change detection without employing directed attention and can be used to index discrimination relevant for language skills (15). In line with the Bayesian Brain Hypothesis (29), MMN represents a prediction error signal (30), reflecting a continuous process in which the brain learns environmental statistics to detect regularity and change, and generates top-down predictions which facilitate stimulus processing (13, 31). The auditory MMN links sensory processing to higher-level cognitive functions and represents processing of violations in the sequence of stimuli, which are compared to the information encoded in the ultra-short-term (echoic) memory (13). Previous source modelling of mMMN has identified bilateral sources of mismatch signal in the superior temporal gyrus and inferior frontal gyrus, reflecting a hierarchical network for processing of prediction error dependent on bidirectional frontotemporal connectivity (32). In the current experiment, the control and ZDHHC9 groups showed deviant-related activity reaching statistical significance in the right temporal cluster, with significant deviant-related responses being more extensive and, mostly, of larger amplitude in the ZDHHC9 group. However, these observations remain qualitative as direct comparison between responses in the two groups was challenging due to limited number of channels for which both groups demonstrated a significant mismatch response. It will be important to replicate these results in larger samples of individuals with *ZDHHC9* variants, within specific age-bands, and in comparison to additional groups such as other monogenic causes of ID.

To explore the network origins of observed between-group differences in MEG signal generation, we applied RNN modelling, in which the activity of hidden units resembles that of neural populations (16, 17). During model training, the RNN weights representing connections between the model units were optimised so that the RNN predictions most closely matched the labels (the grand-average AEF waveforms generated by adding noise to the control group AEF). The feedforward and recurrent weight matrices obtained after training could be interpreted as analogous to wiring patterns supporting neurotypical AEF generation. The RNN model enabled the comparison between its predictions and neurophysiological responses given several analogies that could be made between the two. For instance, creating an RNN model with multiple hidden layers enabled the signal propagation through a hierarchical structure comprising of feedforward and feedback interactions, similar to sensory processing. As the information propagated through the layers, the pattern of activations became more complex, similar to neural signals traveling from sensory periphery to subcortical structures and regions of the sensory cortex. Moreover, the RNN approach facilitated the formation of a high-dimensional activation space given the total of 256 hidden units, which attempted to mimic signals arising from a large number of underlying neural sources that underlie scalp-recorded AEFs.

The RNN model was used as a platform to test a mechanistic hypothesis underpinning AEF generation in the ZDHHC9 group, specifically reduced inhibition arising from ZDHHC9 dysfunction. Inhibitory perturbation of the model after training resulted in increased AEF and deviant-related responses, in keeping with empirical observations although with some limitations. While the model perturbations resulted in an increased synaptic E:I ratio that is smaller than observations in primary cultures with *ZDHHC9* knockout (5), they were appropriate for the current RNN model which captures features of sensory processing, whilst omitting biological details such as separate excitatory and inhibitory units and features of connectivity. Another limitation of the model is that it did not take into account developmental effects. Genetic effects usually interact with the environment continuously, and this interaction shapes sensory processing and behaviours. A future modelling approach would take this aspect into account and might include perturbations from the start of model training as opposed to post-training, potentially in the form of a regularisation term within the loss function equation.

The results of the current study go a step further toward understanding the associations between *ZDHHC9* loss of function, seizure susceptibility and developmental language difficulties. A role for hyperexcitability and atypical AEFs in language development has been proposed in the context of autism, where M100 latencies are persistently delayed and predict language improvement over time (33). M100 latency has been proposed as a marker for capacity to improve cognitive skill, and language skill in particular, relating to cortical interneuron development and GABA concentration (34). Of note, rare variants in GABA receptor subunits have been associated with risk for rolandic epilepsy, potentially bridging seizure risk and cortical inhibition relevant to cognitive development (35). However, it is not known whether M100 amplitude, besides latency, reflects properties of the auditory cortex involved in language acquisition. This could be explored in future prospective studies across monogenic causes of RE, incorporating behavioural measures of speech and auditory processing for correlation with MEG (36). The existing literature provides discrepant accounts of how MMN is affected in epilepsy patients and those affected by developmental language disorders. Some studies revealed links between developmental language disorders and diminished MMN amplitudes, which points to a causal relationship between inefficient auditory processing and insensitivity to phonetic cues and impaired speech and language skills (37, 38, 39, 40). Furthermore, lower MMN amplitude has also been observed in children with rolandic epilepsy, with or without language impairments (41, 42, 43). In contrast, several studies have shown larger MMN responses in epileptic patients in response to pure tones (44, 45, 46). Higher MMN amplitudes in people with epilepsy could indicate increased activation of the same neuronal population as in controls or activation of additional neuronal resources (47). Similar trends have been observed in children with learning difficulties and dyslexia compared to controls (48). These contrasting previous results may reflect small sample sizes and reproducibility issues or could reflect real differences in cortical processing contributing to language difficulties dependent on aetiology and associated network disturbances, which could be explored in future studies.

### Conclusion

In summary, the current study serves as a proof-of-concept for using neural networks to investigate mechanistic origins of developmental cognitive disorders. Future studies will ideally increase the complexity of the neural network models that would better mimic sensory processing and apply these in larger datasets of individuals with cognitive impairments arising from a range of genetic variations. It will also be important to perform studies at more granular scales, for example by studying rodent models of these monogenic disorders and obtaining single-neuron recordings that would shed light on the neuronal dynamics in these conditions. A wider range of cognitive and memory tasks implemented with rodent models and parallel human studies would also be useful to explore effects of single gene variants on learning and memory, together with underlying alterations in neuronal activity and connectivity. Collectively, these inter-disciplinary studies will contribute to improved multi-level understanding of monogenic conditions impacting on cognition and will potentially inform future therapeutic interventions in relevant clinical populations.

## Acknowledgments

We thank the study participants, their families and carers for their extensive contributions and commitment to this project. RIV is funded by a UKRI/MRC PhD studentship. Data collection for this study was funded by the Wellcome Trust/Academy of Medical Sciences (Starter Grant for Clinical Lecturers to KB). This work was supported by UKRI/MRC University Unit Strategic Partnership Funding (programme awards MC_UU_00030/3; MC_UU_00030/2).

## Data availability statement

Data available on request due to privacy/ethical restrictions

## Author contributions

RIV conceived the study, analysed data and wrote the manuscript. JA and DA advised on analyses. DA and KB supervised the project. All authors reviewed, edited and approved the manuscript for submission.

## Ethical statement

Ethical approval for this study was granted by the Cambridge Central Research Ethics Committee (11/0330/EE: Phenotypes in Intellectual Disabilities). Written informed consent was provided by all participants, or a parent / carer for participants under 16 years of age.

