## Supplementary Material for "Synaptic function and sensory processing in ZDHHC9-associated neurodevelopmental disorder: a mechanistic account"

**Figure S1: Auditory evoked neuromagnetic fields (AEF) across all stimuli and corresponding topographic maps**

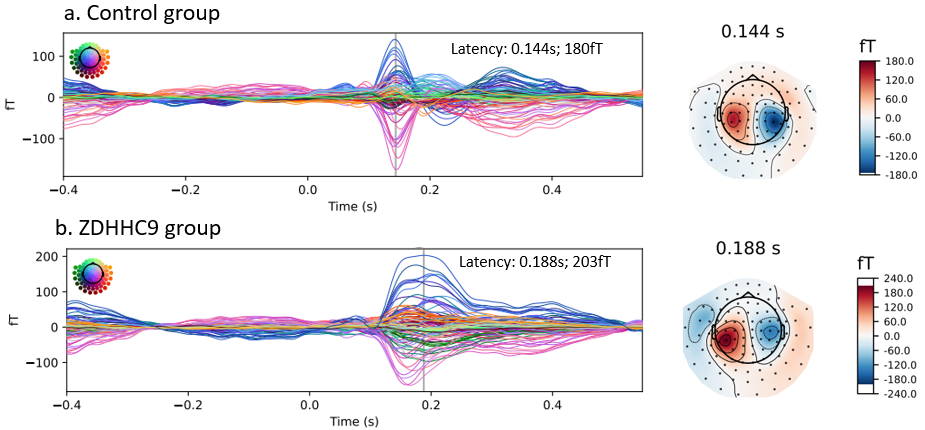

**Figure S2: Frequency-latency dependence of auditory evoked neuromagnetic fields (AEF)**

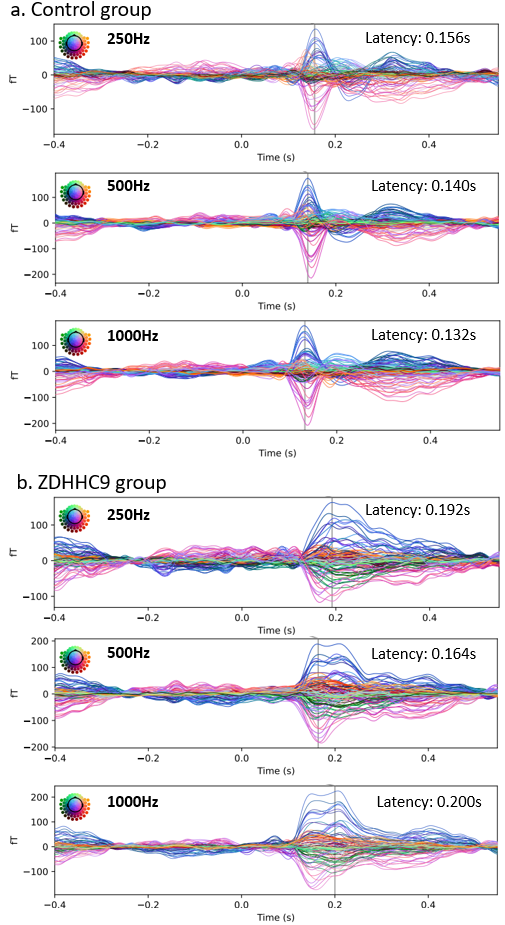

**Figure S3: Direct comparison of mismatch responses in ZDHHC9 and control groups**

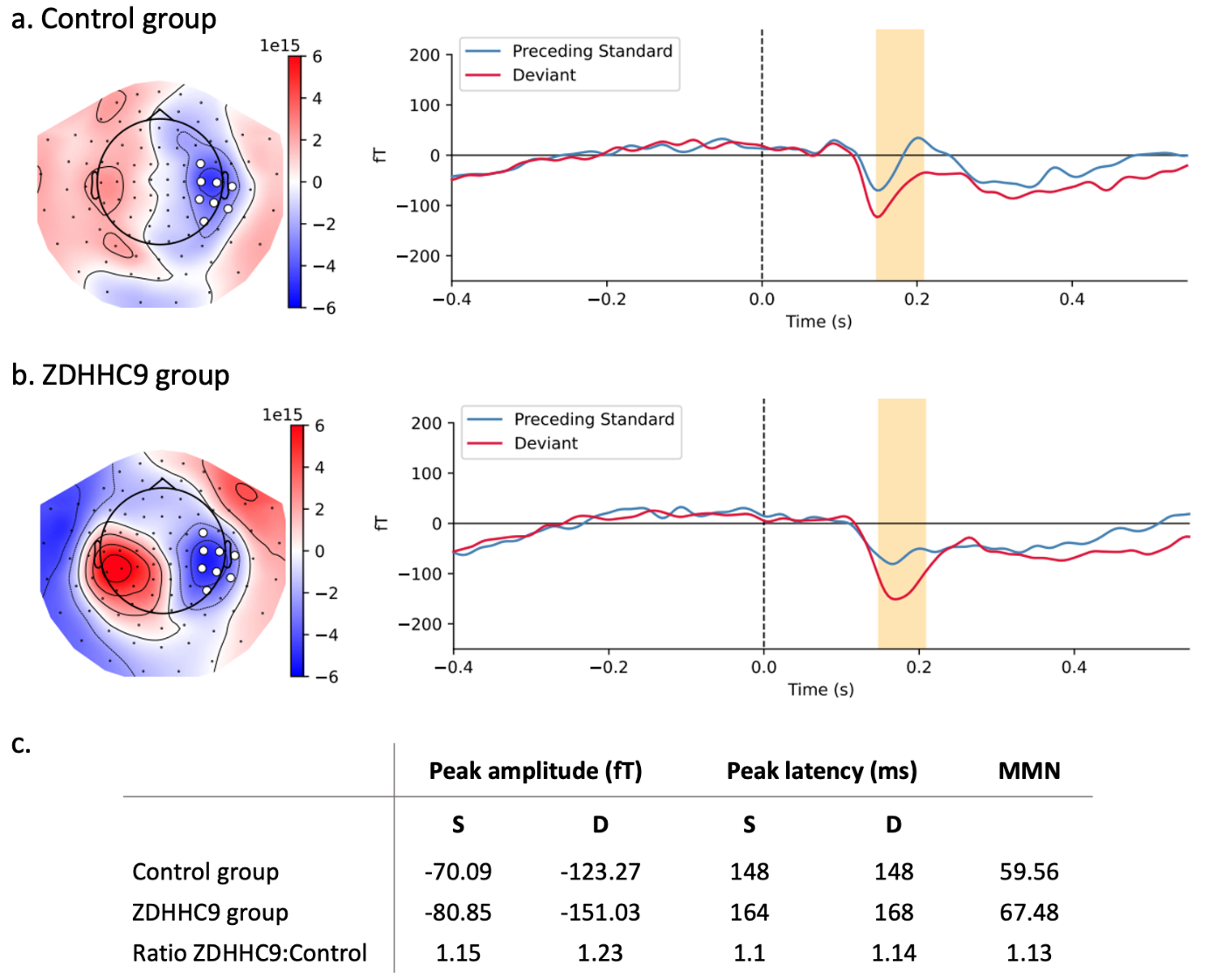

Evoked responses to all deviants (D) and their corresponding preceding standard (S) at the eight sensors where significant S-D differences were found in both the control and ZDHHC9 groups. In yellow, the timeframe where the S-D differences are significant in both groups is shown. **a. C**ontrol group response at the eight overlapping channels of the significant cluster (p-value = 0.0008). **b.** ZDHHC9 group response at the same eight channels of the significant cluster in this group (p-value = 0.0015). **c.** The values from the plots in a. and b. (absolute values for peak amplitudes) and mismatch negativity calculated as mean absolute error between standard-evoked responses and deviant-evoked responses in the significant time window.

**Figure S4: Latent dynamics of 4^th^ hidden layer**

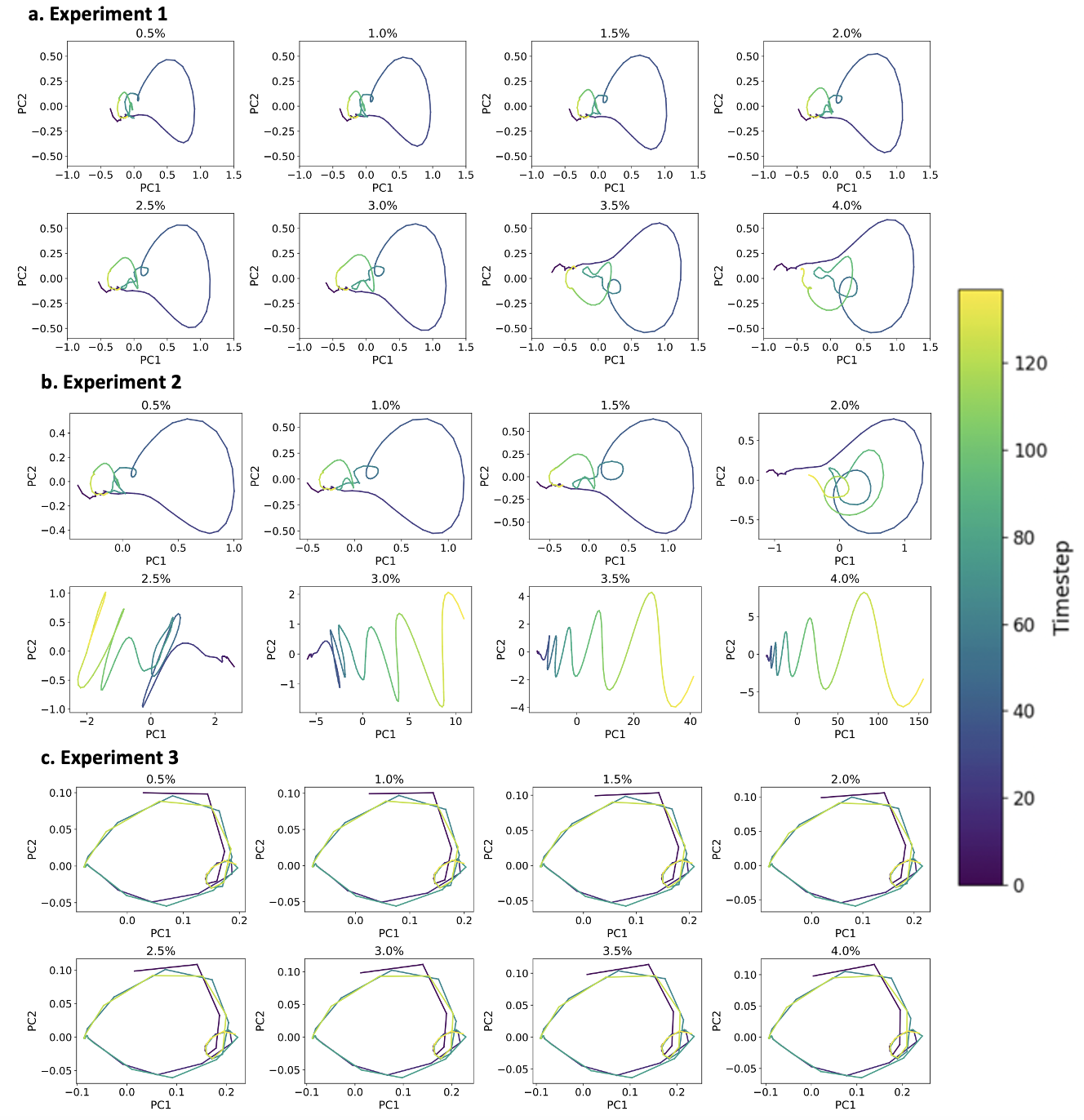

Principal component analysis was performed on the activations of the last hidden layer, which resulted in a latent activity trajectory over time.
